# Extracellular vimentin is sufficient to promote cell attachment, spreading, and motility by a mechanism involving N-acetyl glucosamine-containing structures

**DOI:** 10.1101/2022.11.28.518249

**Authors:** Robert Bucki, Daniel V. Iwamoto, Xuechen Shi, Katherine E. Kerr, Fitzroy J. Byfield, Łukasz Suprewicz, Karol Skłodowski, Julian Sutaria, Paweł Misiak, Agnieszka Wilczewska, Sekar Ramachandran, Aaron Wolfe, Minh-Tri Ho Thanh, Eli Whalen, Alison E. Patteson, Paul A. Janmey

**Affiliations:** Department of Physiology and Institute for Medicine and Engineering, University of Pennsylvania, PA 19104, USA; Department of Medical Microbiology and Nanobiomedical Engineering, Medical University of Białystok, Białystok, Poland; Institute of Chemistry, University of Białystok, Białystok, Poland; Ichor Life Sciences, Inc. LaFayette, NY 13084; Physics Department and BioInspired Institute, Syracuse University, USA

## Abstract

Vimentin intermediate filaments form part of the cytoskeleton of mesenchymal cells, but under pathological conditions often associated with inflammation, vimentin filaments depolymerize as the result of phosphorylation or citrullination, and vimentin oligomers are secreted or released into the extracellular environment. In the extracellular space, vimentin can bind surfaces of other cells and the extracellular matrix, and the interaction between extracellular vimentin and other cell types can trigger changes in cellular functions, such as activation of fibroblasts to a fibrotic phenotype. The mechanism by which extracellular vimentin binds external cell membranes and whether vimentin alone can act as an adhesive anchor for cells is largely uncharacterized. Here, we show that various cell types (normal and vimentin null fibroblasts, mesenchymal stem cells, A549 lung carcinoma cells) attach to and spread on polyacrylamide hydrogel substrates covalently linked to vimentin. Using traction force microscopy and spheroid expansion assays, we characterize how different cell types respond to extracellular vimentin. Cell attachment to and spreading on vimentin-coated surfaces is inhibited by hyaluronic acid (HA) degrading enzymes, HA synthase inhibitors, soluble heparin, or N-acetyl glucosamine, treatments that have little or no effect on the same cell types binding to collagen-coated hydrogels. These studies highlight the effectiveness of substrate-bound vimentin as a ligand for cells and suggest that carbohydrate structures, including the glycocalyx and glycosylated cell surface proteins that contain N-acetyl glucosamine, form a novel class of adhesion receptors for extracellular vimentin.

## INTRODUCTION

Uncontrolled release of intracellular constituents to the extracellular space is generally detrimental to an organism. Potent mechanisms have developed to depolymerize and clear large cytoskeletal filaments such as F-actin from the bloodstream where they otherwise have damaging or pro-inflammatory effects on vasculature and other organs (1–3). Perhaps because of the chemical instability of microtubules in GTP-devoid extracellular fluids no such scavenging system has been described for tubulin, although release of microtubule-binding proteins such as tau is associated with neurodegenerative disease (4–7). Intermediate filaments (IFs), the most stable element of the cytoskeleton, also do not appear to have active extracellular clearance mechanisms, and studies implicate extracellular IF proteins, especially extracellular vimentin, in a range of human diseases (8–12).

Vimentin is a type III intermediate filament protein that forms part of the cytoskeleton of mesenchymal cells, but it can be actively exported to the extracellular environment of cells, and this so-called extracellular vimentin has been shown to perform diverse functions in different settings (9,11,13,14). Its most widely recognized function is that of an intracellular intermediate filament cytoskeletal protein, that modulates the viscoelastic properties of cells as well as spatially regulating various intracellular signals (15–17). Despite not having a signal peptide, or other features of conventionally secreted proteins, vimentin has now been clearly shown to be selectively released into the extracellular space by active processes, rather than appearing in the extracellular environment solely as the result of mechanical damage to the cell membrane permeability barrier. Extracellular expression of vimentin appears to require at least two active processes (18–20). First is activation of protein arginine deaminases or protein kinases that either succinate or phosphorylate key residues that are required for vimentin to assemble into filaments. Activation of these processes leads to solubilization of vimentin into small oligomeric units. The second process needed for vimentin to be released into the extracellular space, or localized to the exterior surface of the cell membrane, involves the unconventional type 3 secretion pathway (21). Once exported to the cell interior or the extracellular space, vimentin can have numerous effects on endogenous host cells or act as an adhesive cofactor for the invasion of specific bacteria or viruses that express on their surface ligands for vimentin (9,11-14).

Many sources of extracellular vimentin that have been identified. When monocytes are activated to macrophages they release vimentin to their exterior membrane surface where it enhances clearance of pathogens and apoptotic cells (8,22,23). Neutrophils release vimentin as a component of neutrophil extracellular traps (NETs) (18) often after citrullination of vimentin (24,25). In wounds to the eye lens that mimic cataract surgery, resident inflammatory cells release vimentin both to their cell surface and to the underlying basement membrane, where it promotes formation of myofibroblasts and a pathologic fibrotic state (26,27). In some tumors, endothelial cells secrete vimentin by the unconventional type 3 secretory pathway, where soluble vimentin activates the VEGF receptor to promote angiogenesis (21). Other cell types release vimentin on the external surface of exosomes where it plays multiple roles in wound healing and development (28,29).

How extracellular vimentin interacts with the eukaryotic cell membranes is less well characterized. Several different transmembrane protein complexes have been shown to be potential receptors for vimentin, but there is not yet consensus for any specific membrane protein as the unique or most important receptor for vimentin. An alternate hypothesis for how vimentin binds the cell surface is based on its affinity for polysaccharides. A compelling study shows that vimentin binds selectively to polymers bearing multiple copies of N-acetylglucosamine, but not most other types of sugar moieties (30). Vimentin also binds glycosaminoglycans, including hyaluronic acid and heparin, in which one of the two carbohydrate units is either N-acetylglucosamine or a closely related structure (31). This affinity for N-acetylglucosamine-containing cell surface structures, and the fact that many or perhaps all of the proposed vimentin binding cell surface proteins are themselves glycosylated, suggest that either the hyaluronic acid- and heparin-rich like glycocalyx or transmembrane proteins that are heavily glycosylated are important elements in the mechanism by which extracellular vimentin binds and activates eukaryotic cells.

Whether vimentin in a basement membrane or a substrate is sufficient to promote cell adhesion or whether it primarily acts to modify adhesion to more common extracellular matrix (ECM) ligands such as fibronectin or collagen is also unknown. In this study, we prepare inert hydrogel substrates that bear only covalently attached vimentin as a potential cell ligand and show that multiple cell types bind these surfaces, which promote spreading and motility, but not stress fiber formation, large focal adhesion formation, or cell proliferation.

## EXPERIMENTAL PROCEDURES

### Reagents

D-glucuronic acid (G526), 4-methylmubelliferone (M1381), hyaluronic acid sodium salt from *Streptococcus equi* mol wt 15,000-30,000 (97616), 1,6-Hexanediol (H11807), N-Acetyl-D-glucosamine (A3286), heparin sodium salt from porcine intestinal mucosa (H3393), D-Mannitol (M4125), (3-Aminopropyl)trimethoxysilane (281778) and 70% glutaraldehyde (G7776) were from Sigma-Aldrich, St. Louis, MO, USA. 2% bis acrylamide (1610142) and 40% acrylamide solutions (1610140) were from Bio-Rad Laboratories, Hercules, CA. TEMED (TB0508) was from Bio Basic. Ammonium persulfate (17-1311-01) was from Life Sciences, Uppsala Sweden, hyaluronidase from *Streptomyces hyaluronlyticus* (389561) was from Millipore Corp., USA. Collagen I was from BD Bioscience, San Diego, CA. Sulfo-SANPAH cross-linker (A35395) was from Thermo Fisher Scientific. Recombinant human protein arginine deiminase 4 (PAD4) was from Cayman Chemical, Inc.

### Preparation of soluble vimentin

Vimentin-coding DNA was cloned into the pET7 plasmid (provided by Robert Goldman, Northwestern University), transformed into Escherichia coli BL21(DE3) competent cells and grown on LB-Agar plates containing 100 mg/mL ampicillin. Single colonies from the LB-Agar plates were inoculated into LB medium containing 100 mg/mL ampicillin and grown overnight at 37°C. The overnight cultures were used to inoculate a 10 L bioreactor containing LB medium at 37°C that was agitated at 600 RPM. Protein production was induced by the addition of IPTG to a final concentration of 100 mM, and the cultures were left overnight in the bioreactor. Cells were harvested and resuspended in lysis buffer (50 mM Tris-HCl (pH 8.5), 0.5 M NaCl, benzonase and lysozyme). The resuspended cells were lysed by sonication, and the suspension was centrifuged on a JS-4.2 rotor (Beckman) at 6000 RPM for 30 minutes. The pellet was washed successively in wash buffer-1 (50 mM Tris-HCl (pH 8.5), 1 M NaCl, 0.1% Triton X-100, 2 M urea) and wash buffer-2 ((50 mM Tris-HCl (pH 8.5), 1 M NaCl, 2 M urea) by resuspension and centrifugation on a JS-4.2 rotor (Beckman) at 6,000 RPM for 30 minutes. The washed pellet was resuspended in solubilization buffer (50 mM Tris-HCl (pH 8.5), 8 M urea) by stirring overnight at room temperature. The solubilized pellet was loaded on a HiTrap Q FF column (5 mL volume; Cytiva) that had been equilibrated in solubilization buffer and eluted in the same buffer using a 0 to 1 M gradient of NaCl. Fractions containing vimentin were pooled and snap-frozen in liquid nitrogen. The purity of vimentin was assessed using SDS-PAGE (S1 Fig 1). A fraction of the vimentin preparation was labeled with rhodamine B-succinimide by the method previously used to label neurofilaments (32).

### Citrullination of vimentin

The citrullination procedure was based on a method for citrullinating fibrinogen (33). Purified vimentin (4 mg/ml) was dialyzed into 10 mM Tris-HCl, 1 mM EDTA and 6 mM DTT, pH 7.6, a low ionic strength buffer that destabilizes vimentin polymerization (V1). A portion of vimentin was supplemented with 2 mM MgCl2 and 150 mM KCl to polymerize it (V2). Lyophilized PAD4 was suspended in 40 mM Tris-HCl, 5 mM CaCl2 and 10 mM DTT, pH 7.5 to a concentration of 3 mg/ml. Vimentin and PAD4 were mixed to concentrations of 3.8 mg/ml and 0.13 mg/ml, respectively and incubated 12 hrs at 37°C. Purity of the vimentin preparations and evidence of its citrullination are shown in SI Fig. 1. As reported previously (34), vimentin citrullination results in filament disassembly under polymerizing conditions.

### Preparation of vimentin- collagen- and fibronectin-coated substrates

Cells were cultured on polyacrylamide hydrogel substrates of 0.5 and 30 kPa stiffness that were prepared using the methods previously described (35,36). The acrylamide and bis-acrylamide solutions were formulated in distilled H_2_O to a total volume of 1 mL. 1 μL of TEMED and 3 μL of 2% ammonium persulfate were used as polymerization initiators. The solution (30 μL) was deposited on a 12-mm square glass coverslip pretreated with 3-aminopropyltrimethoxysilane and 0.5% glutaraldehyde. After removing the top coverslip, polyacrylamide gels were covalently linked to ligands by incubating the gels with 50 μL of 0.1 mg/mL collagen I, 0.1 mg/mL fibronectin or 0.1 mg/mL vimentin after activating the gel surface with the UV-sensitive Sulfo-SANPAH cross-linker. The stiffness of the gel substrates was verified using atomic force microscopy (data not shown).

### Cell types and cell culture

Wild-type and vimentin-null mouse embryonic fibroblasts provided by J. Ericsson (Abo Akademi University, Finland) were maintained in Dulbecco’s Modified Eagle’s Medium + 4.5g/l glucose + 2mM L-glutamine + sodium pyruvate (Corning) supplemented with 10% fetal bovine serum (Hyclone), 1% nonessential amino acid (Fisher Scientific), 25 mM HEPES (Fisher Scientific), 100 μg/mL streptomycin, and 100 units/mL penicillin (Fisher Scientific). Cell cultures were maintained at 37°C with 5% CO_2_. Cells were passaged every 3–4 d and harvested for experiments when 70% confluent. The lung carcinoma cell line A549 CCL-185 (ATCC) was cultured in DMEM (Gibco) supplemented with 10% fetal bovine serum (FBS, Gibco, Grand Island, NY) 100 U/mL penicillin and 100 μM streptomycin (Sigma-Aldrich, St. Louis, MO) on tissue culture plastic and kept in a humidified incubator at 37 °C and 5% CO_2_. Human mesenchymal stem cells (hMSCs) (Lonza) were cultured in the same conditions in their respective medium for a period of 24 hours or greater.

### Adhesion, motility, proliferation studies by live cell imaging

Cell area, circularity, and proliferation were determined from optical images collected with 10x or 40× air lens and phase contrast using a Hamamatsu camera on a Leica DMIRE2 inverted microscope (Leica). For each condition, a minimum of 5 (10X) – 20 (40x) images was acquired, and approximately 100 – 150 cells per condition were analyzed. Cell area was calculated using Fiji software by tracing cell peripheries (Fiji Software, NIH, Bethesda, MD). Only single cells were taken into account. The migration speed of cells was determined with time-lapse microscopy over a period of at least 10 h in 5 min intervals. A Tokai-Hit Imaging Chamber (Tokai Hit, Shizuokaken, Japan) that maintained a humid 37 °C and 5% CO_2_ environment was first equilibrated for 1 h. Cell cultures were placed inside the chamber mounted on a Leica DMIRE2 inverted microscope (Leica) equipped with an ASI *x/y/z* stage (BioVision Technologies) and a Hamamatsu camera; a 10X air lens was used for image sequence recording. Cell migration speed *u* (length of the total trajectory *d* divided by time *t*) was calculated by tracing the (*x,y*) position of the center of the cell nucleus at every image using ImageJ Software (NIH) and the Manual Tracking plugin (https://imagej.nih.gov/ij/).

### Immunofluorescence

Cells were fixed with 4% paraformaldehyde (Sigma) for 10 min at room temperature, permeabilized with 0.1% Triton X-100 in TBS for 10 min, and then incubated for 1 h with primary antibodies directed against vinculin (1:1000; mouse monoclonal anti-vinculin IgG; sc-73614 Santa Cruz). In the next step the cells were incubated with secondary antibodies: 1:1000; Alexa-Fluo 568 goat anti-mouse IgG; A-11004 (Invitrogen), phalloidin–tetramethylrhodamine B isothiocyanate (1 μg/ml; Sigma) and DAPI (1 μg/ml; Sigma).

### Traction force microscopy

Cell traction force was determined by first measuring the hydrogel displacement field and then solving the inverse elastic problem to reconstitute the traction force map. Specifically, red fluorescent beads (200 nm, Thermo Fisher Scientific) were mixed at a density of 0.3% v/v with the 30 kPa polyacrylamide pre-gel solution containing 10.4% acrylamide and 0.264% bis-acrylamide. After polymerization, hydrogels were activated with 0.2 mg/ml Sulfo-SANPAH and then coated with 0.1 mg/ml vimentin (36). Human MSC cells were sparsely seeded on the vimentin-coated hydrogels to minimize the cellcell contact. A bright field image of a randomly selected cell and the corresponding fluorescent beads image was taken. The cell was then detached from the hydrogel with 0.5 N sodium hydroxide, and the reference image of the fluorescent beads was taken. The in-plane displacement field of the hydrogel substrate was calculated by particle image velocimetry. The traction force field was then constructed by solving the inverse problem of Boussinesq solution through Fourier transform traction force cytometry (37,38).

### Dynamic light scattering

Dynamic light scattering (DLS) was used to assess the binding of soluble vimentin to potential ligands that could affect its binding to the cell surface. The average aggregate size (hydrodynamic diameter) was determined using a Zetasizer Ultra (Malvern Panalytical Ltd, UK). The method measures the diffusion constant of solutes from the autocorrelation function of scattered light intensity. The diameter is calculated from the relation D = kT/6πηRh, where D is the translational diffusion constant, η is the solvent viscosity, and Rh is the hydrodynamic radius. 40 μl samples of vimentin (at 0.1 mg/ml), suspended in PBS were placed in cuvettes and then incubated with GlcNAC, 1,6-hexanediol, heparin and D-mannitol (0.05 mg/ml) for 5 min before data acquisition.

### 3D Spheroid formation and analysis

To perform cell aggregate invasion assays, cell aggregates were first generated using the procedure described by Ibidi® (39). Briefly, 96-well flat-bottom plates were coated with 40 μl of 1% agarose solution diluted with PBS. Wild-type and vimentin-null mEF were then plated at 30000 cells/well in 100 μl of DMEM/F12 (10% FBS, 1xPen/Strep). Cell aggregates were incubated at 37°C with 5% CO_2_ for 2 days before harvesting. The diameter of the spheroids was between 100 to 200 μm when collected. Collagen gels were prepared using a slightly modified protocol from Ibidi® (40). Collagen type I, rat tail 5mg/ml (Ibidi) were diluted on ice to 1.5mg/ml with 6.67% of 5x DMEM (Fisher scientific), 10% FBS in 1x DMEM. The pH was adjusted to 7.4 using 1M NaOH and sodium bicarbonate. Cell aggregates were added to the collagen solution before it solidified. The collagen solution is then pipetted on 1% agarose coated 24-well flat bottom plates and let polymerized for 40 minutes in incubator at 37°C, 5% CO_2_ and 100% humidity. To assess the aggregates’ invasion through the collagen network alone or with the presence of either native vimentin or citrullinated vimentin, 2 μg/ml of each type of vimentin was added before polymerization. DMEM complete media was added on top of the fully polymerized collagen gel. After 3 hours inside the incubator, spheroids in collagen gel are imaged for 48 hours with phase -contrast widefield using a Plan Fluor 10× objective on Nikon Eclipse Ti inverted microscope equipped with an Andor Technologies iXon em+ EMCCD camera (Andor Technologies). Aggregates were maintained at 37 °C and 5% CO_2_ using a Tokai Hit (Tokai-Hit) stage top incubator. Cross-sectional area of the spheroid (core and connecting strands) was traced using ImageJ.

## RESULTS

### Adhesion to vimentin-coated gels

To test whether substrate-bound vimentin was sufficient for cell adhesion, we prepared inert polyacrylamide (PAAm) hydrogels, which resist cell adhesion, and covalently laminated their surface with vimentin, fibronectin, or collagen I (Fig. 1). Figure 1a shows a schematic of the protein-coated gel substrate. The purity of the bacterially expresses vimentin is shown in SI Fig 1. Previous studies have shown that surface treatment of these gels with an integrin ligand or other cell attachment factor is necessary for cells to form stable adhesions and survive on these surfaces. To confirm binding of vimentin to the polyacrylamide gel surface, we stained either collagen-coated (Fig. 1b) or vimentin coated (Fig. 1c) PAAm gels with anti-vimentin antibodies and detected vimentin only on the surface of the gel vimentin-coated gel with immunofluorescence imaging. The surface coating of these gels is relatively uniform, but also shows some submicron scale texture due to the polymeric state of the vimentin and the imperfect reaction of the linker group with the polyacrylamide gel surface. These gels were also used to detect possible binding of soluble vimentin to a collagen-coated substrate. Figures d,e show that rhodamine-labeled vimentin does not bind directly to an uncoated polyacrylamide gel (1d), but it binds to gels that are covalently laminated with collagen-I (e).

**Figure 1.**
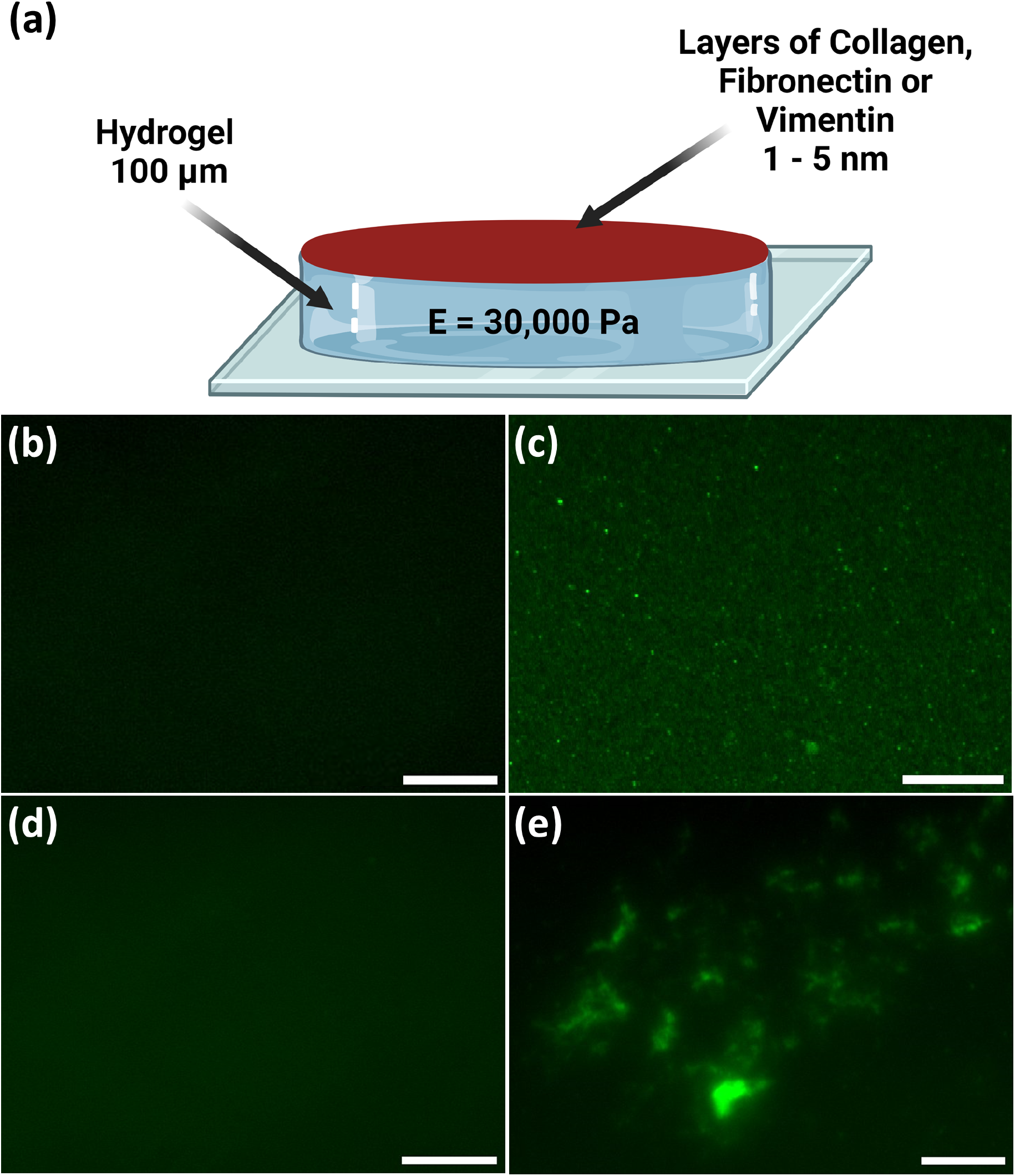
**(a)** Scheme of polyacrylamide gel coated with different proteins. 30 kPa gel with collagen **(b)** or vimentin **(c)** stained with anti-vimentin antibody. 30 kPa uncoated **(d)**, or collagen-coated **(e)** gel stained with rhodamine-labeled vimentin. Scale bar, 20 μm.

Figure 2 shows that human mesenchymal stem cells and A549 lung cancer cells adhere and spread on vimentin-coated polyacrylamide gels, suggesting that adhesion to vimentin-containing substrates might be a common feature of cells. Both the kinetics and extent of cell spreading on vimentin-coated gels are different from the spreading rate on collagen-coated gels or glass, suggesting that the receptors and downstream signals activated by engagement of vimentin might be distinct from those that govern cell adhesion to integrin ligands.

**Figure 2.**
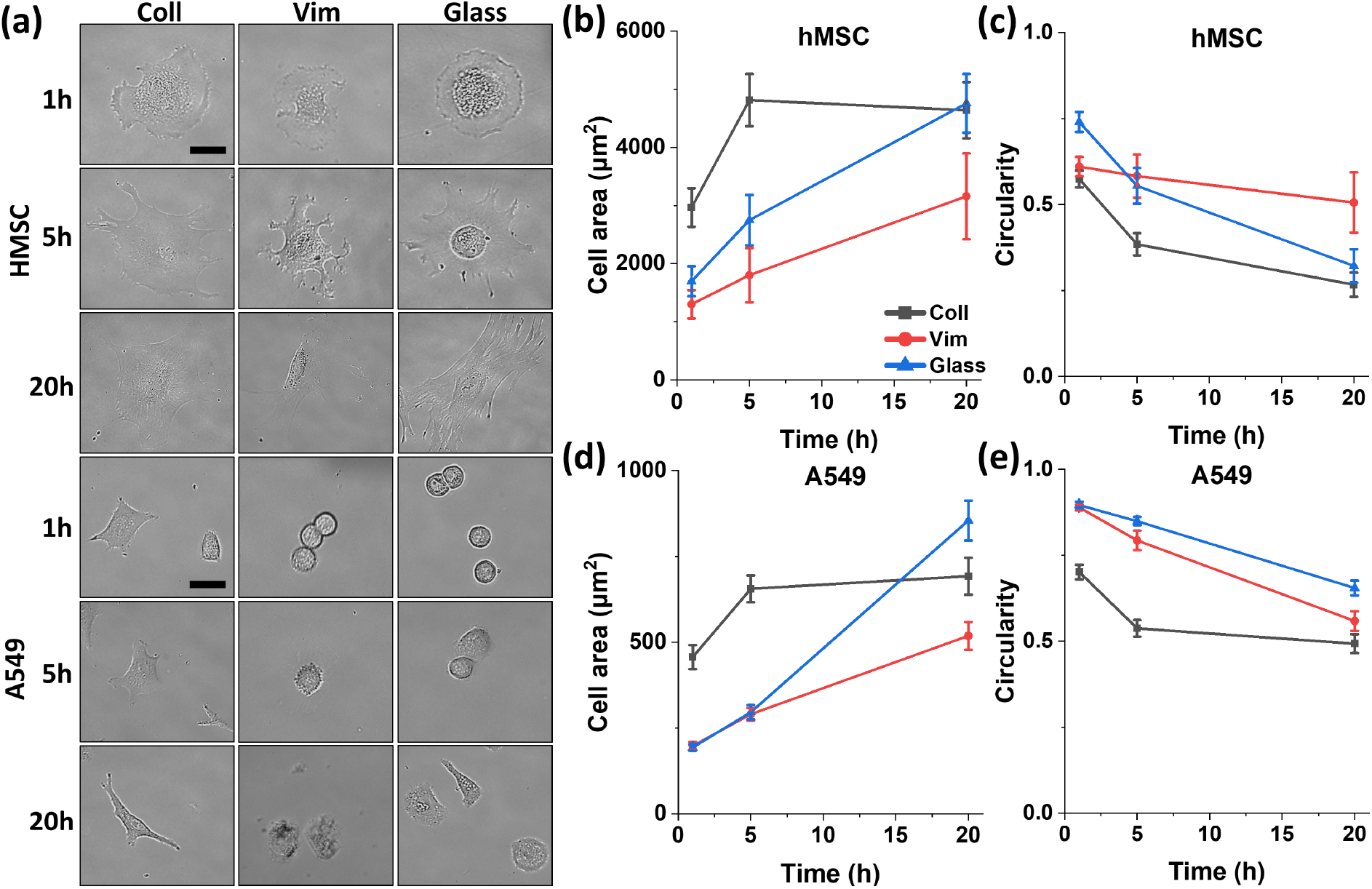
Morphology and kinetics of spreading of hMSCs and A549 cells on glass or 30 kPa PAAm gels coated with collagen or vimentin. **(a)** Live cell images of normal or vimentin null mEFs after 16 hrs on 30 kPa vimentin-coated PAAm gels. **(b-e)** Time course of changes in spread area and circularity of hMSCs and A549 cells on glass or 30 kPa PAAm gels coated with collagen or vimentin. Scale bar, 20 μm.

Figure 3 shows that both normal and vimentin null mouse embryo fibroblasts (mEF) adhere to vimentin-coated 30 kPa PAAm gels and spread on them to form flattened cell structures, similar in size to those they would form on integrin ligand coated gels or glass or plastic surfaces. The similar morphologies of normal and vimentin-null fibroblasts on vimentin-coated substrates show that adhesion of these cells to substrate-bound vimentin does not require vimentin-vimentin contacts, which might occur in vimentin positive cells that also express vimentin on their surface. Therefore, the vimentin adhesion receptor on these cells appears to be distinct from vimentin itself and might function either similarly or differently from other cell adhesion proteins, such as integrins. Fibroblasts bound to vimentin-coated gels are capable of mechanosensing, since the spread area of cells on 5 kPa vimentin-coated gels was much smaller than those on 30 kPa gels (SI Fig 2). Lack of integrin activation by vimentin-coated gels is also supported by the observation that no cell division was seen over 20 hours on vimentin-coated gels, whereas 5% of the fibroblasts divided on collagen- or fibronectin-coated gels.

**Figure 3.**
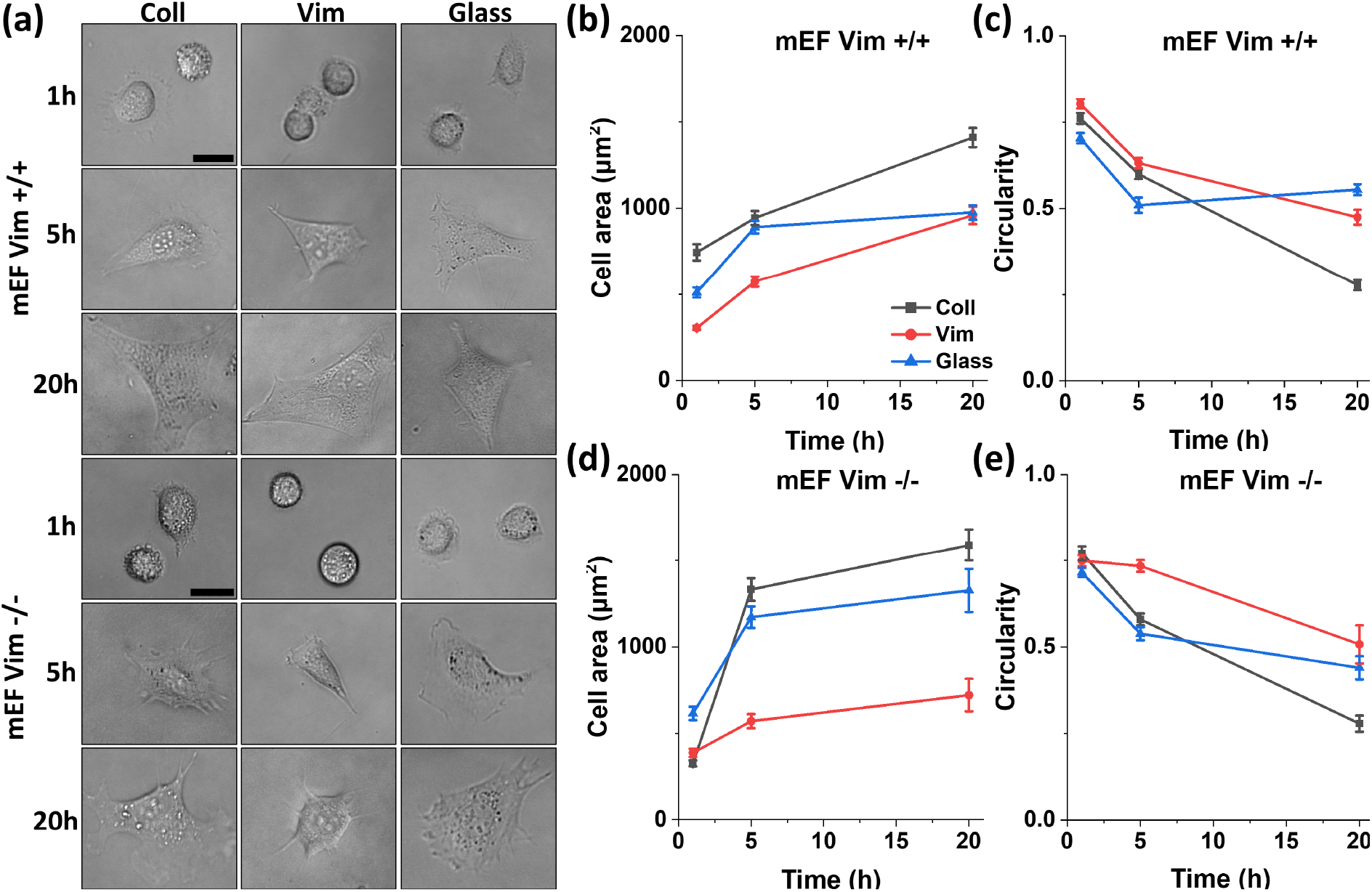
Morphology and kinetics of spreading of vim +/+ and vim -/- mEFs on glass or 30 kPa PAAm gels coated with collagen or vimentin. **(a)** Live cell images of normal or vimentin null mEFs after 16 hrs on 30 kPa vimentin-coated PAAm gels. **(b-e)** Time course of changes in spread area and circulatory of vim +/+ and vim -/- mEFs on glass or 30 kPa PAAm gels coated with collagen or vimentin. Scale bar, 20 μm.

### Cytoskeletal changes and adhesion sites in vimentin substrate-bound cells

Consistent with the hypothesis that cell adhesion to vimentin-coated surfaces is distinct from adhesion to collagen or other integrin ligands, Figure 4 shows that although cells spread on both collagen-coated and vimentin-coated surfaces, the structure of their actin and vimentin filament networks is different. Similarly, the adhesion sites that these cells form are different. Figure 4a-c shows that the formation of large actin bundles, or stress fibers, is a common feature of fibroblasts and MSCs binding to stiff collagen-coated gels, but these structures do not appear, or at least are much less developed, in cells spread to similar areas on vimentin-coated gels (Figure 4d-f). Vinculin is more localized to actin-containing fibers and less diffuse for cells on collagen (a-c) compared to cells on vimentin (d-f). A549 lung cancer cells appear to be different. They form actin bundles on both collagen-coated (c) and vimentin-coated gels (e), although the vinculin staining is less diffuse on collagen-coated gels than vimentin-coated ones (e).

**Figure 4.**
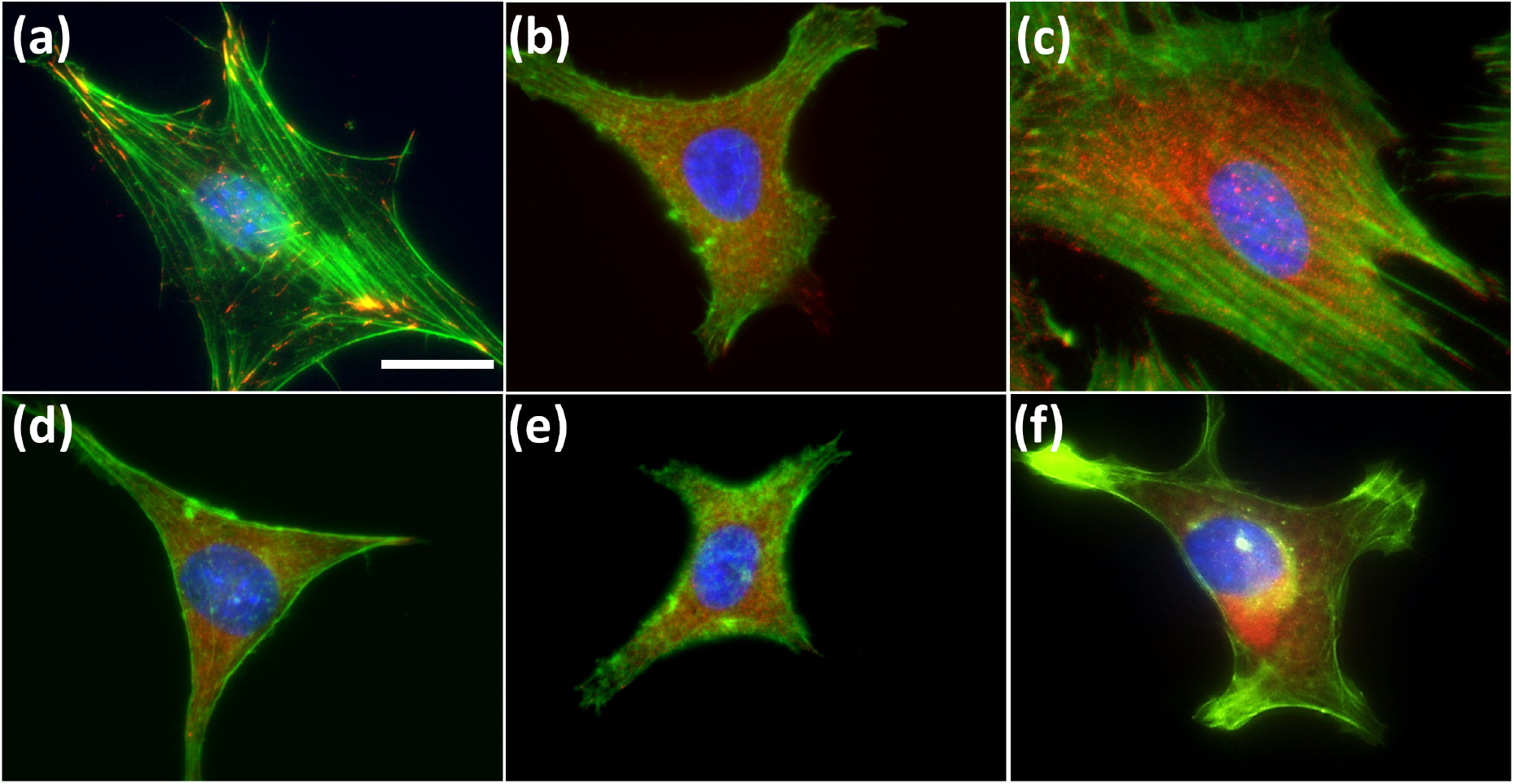
Fluorescence images of mEF Vim +/+ **(a, d)**, A549 **(b, e)** and HMSC **(c, f)** cells seeded on 30 kPa hydrogels coated with collagen **(a-c)** or vimentin **(d-f)**. Vinculin is presented in red, F-actin in green. Cells were counterstained with DAPI (blue). Scale bar, 20 μm.

### Traction stresses on vimentin-coated deformable substrates

Despite the lack of actin stress fibers or large focal adhesions in cells bound to vimentin-coated gels, these cells nevertheless are able to exert traction stresses on the substrate, consistent with their ability to mechanosense (SI Fig 2). Figure 5a shows a typical MSC bound to a vimentin-coated gel and a typical strain field generated by the cells on the deformable substrate using traction force microscopy. Figure 5b shows the average spread area on gels coated with collagen, fibronectin or vimentin, and Figure 5c shows that the magnitude of traction stress that cells exert on vimentin-coated surfaces is significant, but smaller than the traction stresses on collagen- or fibronectin-coated substrates. These results are consistent with the morphology shown in Figures 3-4 and confirm that the nature of the adhesion of cells to vimentin-coated substrates is different from that of integrin-dependent adhesion.

**Figure 5.**
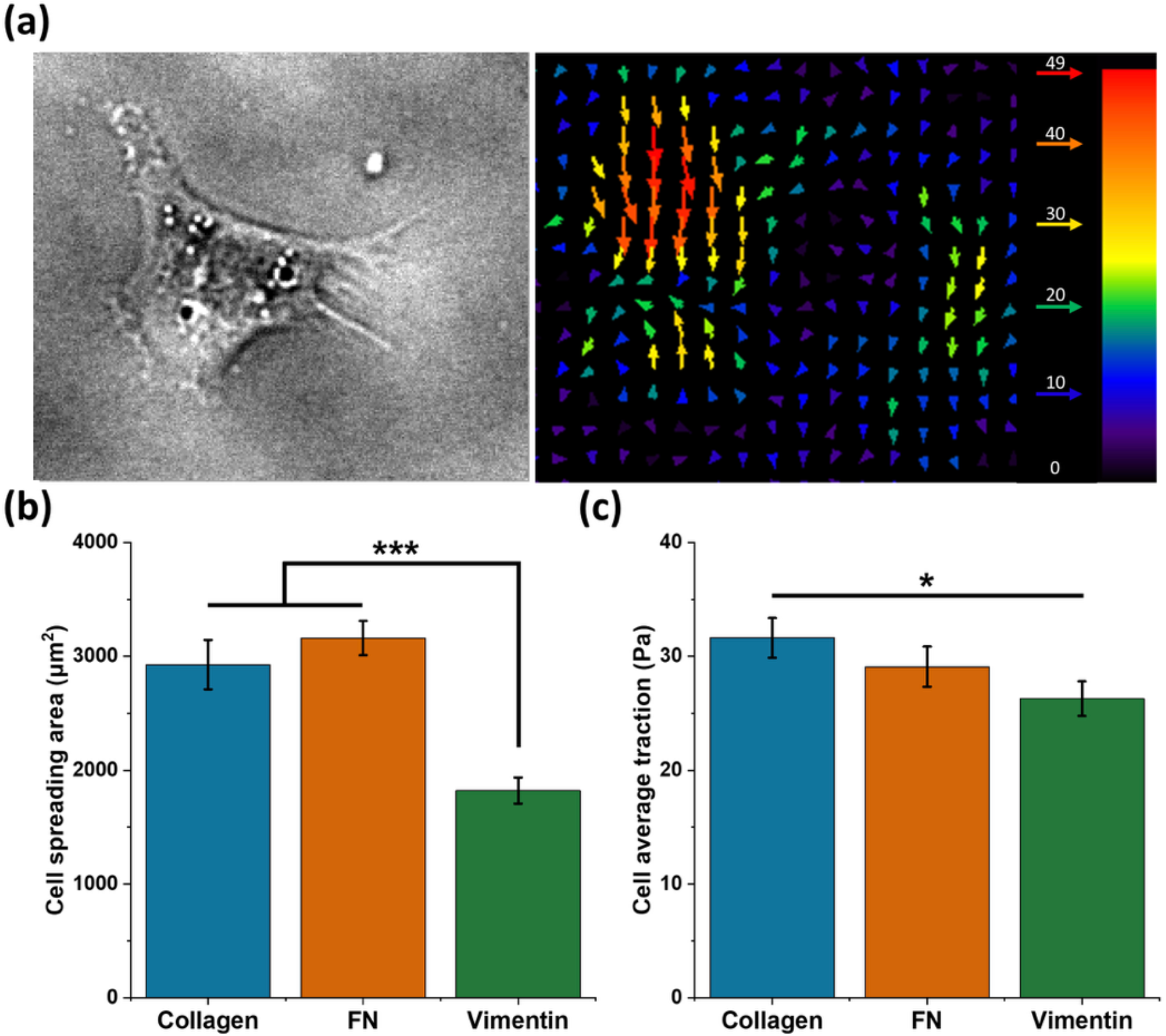
Traction stress measurements of hMSCs on 30 kPa PAA gels coated with collagen, fibronectin, or vimentin.

### Extracellular vimentin promotes cell migration through ECM networks

Previous studies suggest that citrullinated vimentin is more effective than unmodified vimentin in promoting motility of cells through extracellular matrices (41). To determine whether extracellular vimentin mediates cell motility through three-dimensional (3D) fibrillar collagen matrices, we prepared spheroidal aggregates of mEF embedded in 2 mg/mL collagen I networks with the addition of either vimentin or its citrullinated form (2 ug/mL) to the collagen network (Fig. 6). Vimentin binds collagen as indicated by Fig 1, and immunofluorescence staining showed vimentin was incorporated into the collagen network. We found that in the addition of extracellular vimentin (in contrast to its citrullinated form) did not change the average spheroidal aggregate area over time but did impact how cells migrated into the collagen gel, which was individually rather than as strands of connected cells (Fig. 6 h,i). Using time-lapse imaging of the embedded cell aggregates, we observed that invading strands of cells do form in the presence of extracellular vimentin but many of them disassemble over the course of 24 hours, similar to the effect of vimentin on endothelial cells (21). In contrast to normal vimentin in the network, citrullinated vimentin significantly increased collective invasion of wild-type mEF into the collagen network, as shown in Fig. 6. This is consistent with the effect of citrullinated vimentin on primary human lung fibroblast migration (41), and here we note markedly large strands of cells that emerge from the aggregate in the presence of the additional citrullinated vimentin in the network (SI Fig. 3). The invasion of spheroidal aggregates formed of vimentin-null mEF into the collagen network is similar to that of aggregates formed of wild-type mEF. Addition of full vimentin and citrullinated vimentin to the collagen network did not increase vimentin-null aggregate spread areas but did increase the number of individual cells that escape into the network.

**Figure 6.**
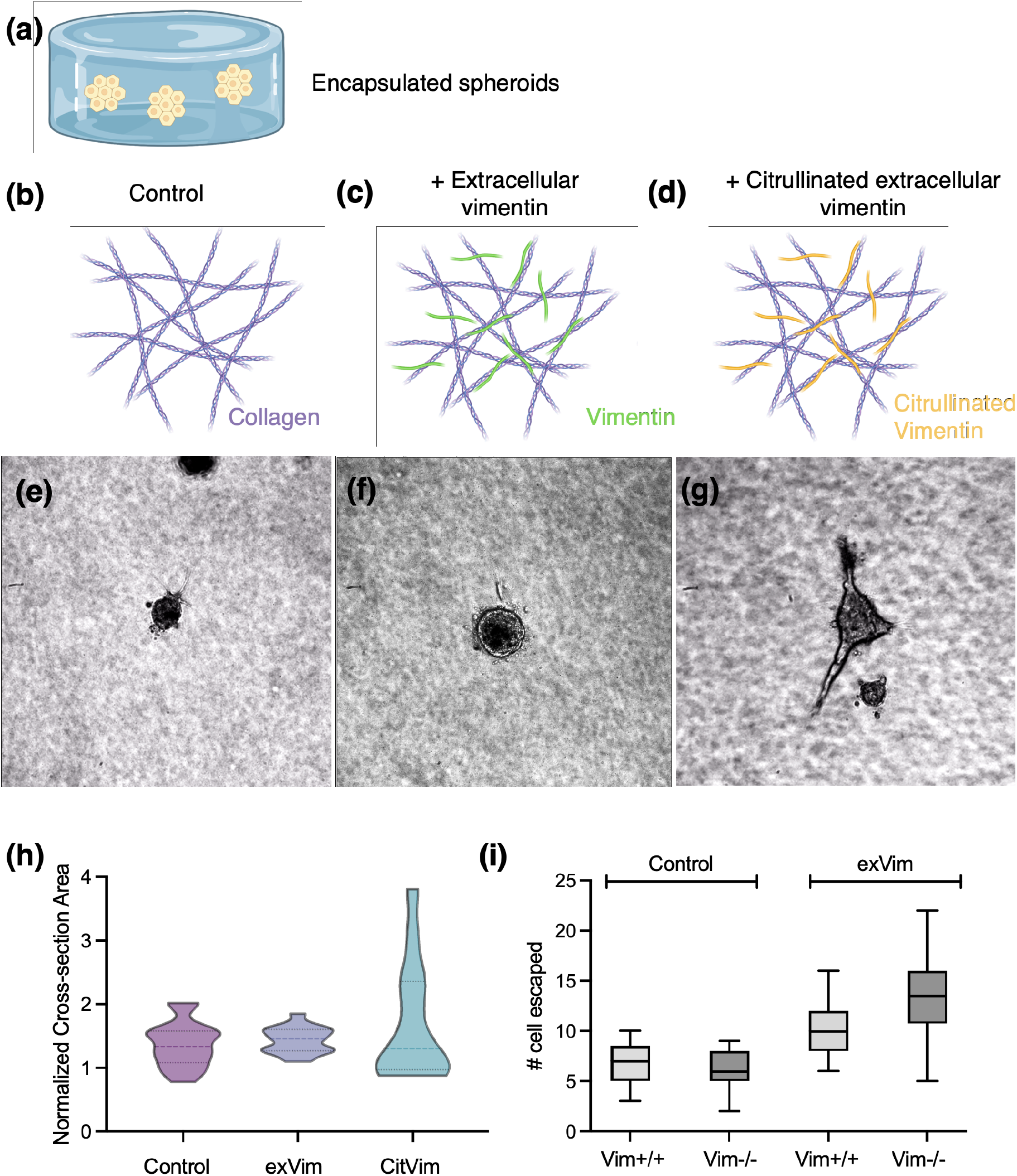
Extracellular vimentin promotes cell migration through ECM networks. (a) Experimental method. Cellular aggregates are embedded in collagen I gels (2 mg/mL). Cartoon representing the matrix presentations to the cells: (b) the control collagen I matrix, (c) collagen I with extracellular vimentin (2ug/mL), and (d) collagen I with extracellular vimentin in its citrullinated form (2ug/mL). Representative phase-contrast images of cell aggregates formed by wild-type mEF embedded in matrixes containing (e) collagen I, (f) collagen I + vimentin, and (g) collagen I + citrullinated vimentin at 24 hours. (h) Averaged projected area of cell aggregates formed from wild-type mEF embedded in collagen network (24 hr). (i) Single cell escape data for wild-type and vimentin-null MEF aggregates for each condition (collagen, collagen+vimentin, collagen+ citrullinated vimentin). Data collected at 48 hr. Addition of extracellular vimentin to the network increases number of cells escaping from aggregates for both cell types.

### Effects of glycocalyx and N-acetyl glucosamine on adhesion to vimentin-coated substrates

Based on previous reports that vimentin can bind hyaluronic acid (42) and heparin (43), constituents of the cells’ glycocalyx, we tested the effect of hyaluronidase, which degrades the glycocalyx, and the drug 4’MU, which inhibits synthesis of the glycocalyx by the membrane bound enzyme HA synthase. Figure 7a shows that hyaluronidase treatment strongly decreases the adhesion of fibroblasts and MSCs to vimentin-coated substrates, which has no effect on the adhesion of the same cell types the collagen-coated gels. Figure 7b shows that cells that do attach to vimentin-coated gels after inhibition of glycocalyx production, have significantly smaller spread area, but the effect that is not statistically significant for cells on collagen-coated gels. In addition, we tested the effect of hyaluronidase treatment on the motility of cells on collagen- or vimentin-coated substrates. Figure 7c shows that the motility of hMSCs was almost completely blocked by hyaluronidase, whereas there was a much smaller effect on the motility of cells on collagen-coated shells, an effect that is expected because of the interplay between the glycocalyx and integrins (44).

**Figure 7.**
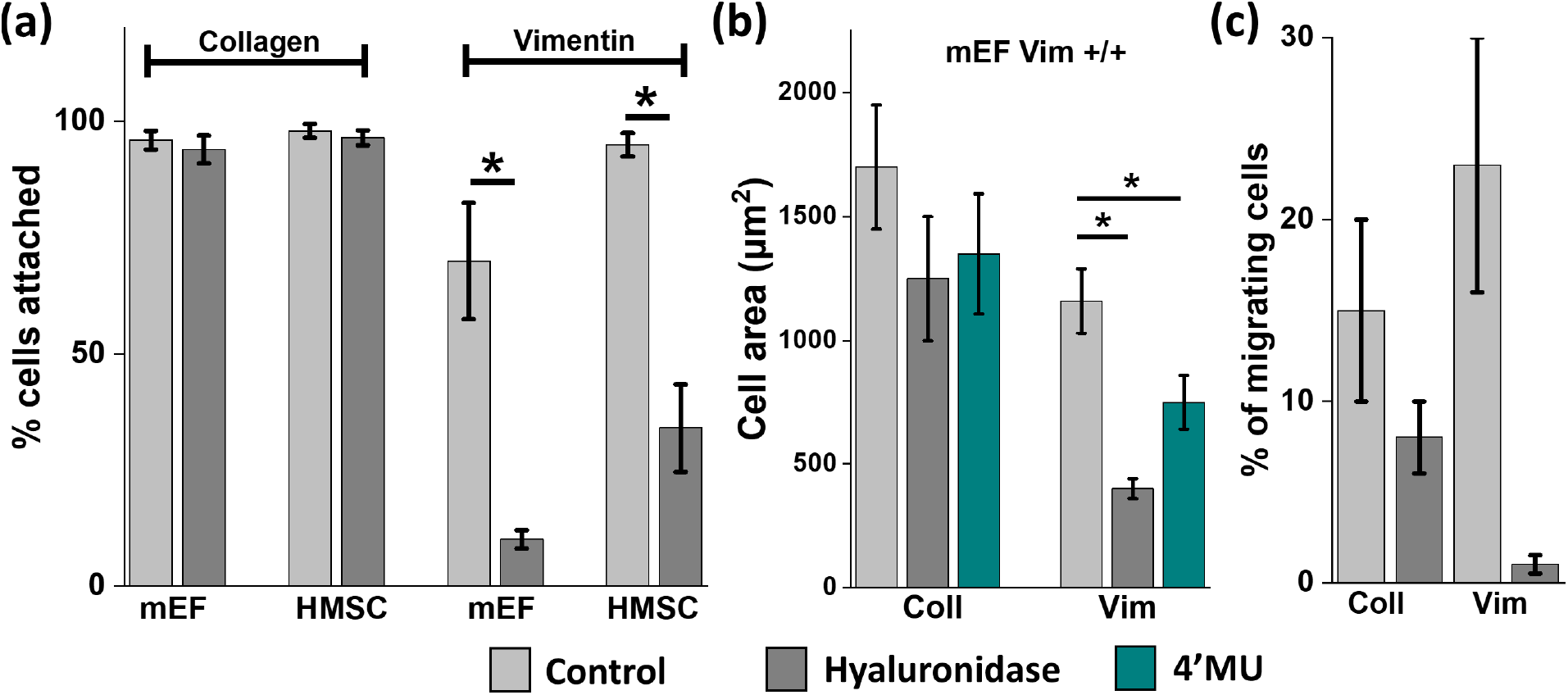
**a)** Fibroblast and hMSC attachment is inhibited by hyaluronidase on vimentin-coated but not collagen-coated PAA gels. **b)** Average cell area of mEFs cultured on PAA gels coated with collagen (left) or vimentin (right) +/- hyaluronidase or 4’MU. **c)** hMSC migration on vimentin-coated gels is inhibited by hyaluronidase more than on collagen-coated gels.

Previous studies have shown that cells are able to bind otherwise nonadhesive surfaces that are coated with polymers containing N-acetylglucosamine (GlcNAc) but not other sugars, and that the receptor on the cell surface, which acts like a lectin, is vimentin (45). Cell adhesion in this study was blocked if the cells were treated with soluble GlcNAc before exposure to the GlcNAc polymer-containing substrate. We therefore tested the effect of soluble GlcNAc on cell adhesion to vimentin-coated surfaces. Figure 8 shows that soluble GlcNAc strongly inhibits the adhesion to and spreading of fibroblasts on vimentin-coated surfaces, but it has no effect on the adhesion of the same cell type to collagen-coated gels. The glycosaminoglycan heparin, which contains GlcNAc subunits also strongly inhibits binding of cells to vimentin-coated gels, but has no effect on collagen-coated gel adhesion. In contrast, 1-6 hexane diol, which has been reported to interfere with vimentin-vimentin interactions (46), has no effect on adhesion of cells to either vimentin- or collagen-coated substrates, confirming that cell adhesion to vimentin-coated surfaces does not involve vimentin-vimentin contact. D-mannitol, previously shown not to affect binding of vimentin-coated cells to GlcNAc-coated surfaces (45), also had no effect.

**Figure 8.**
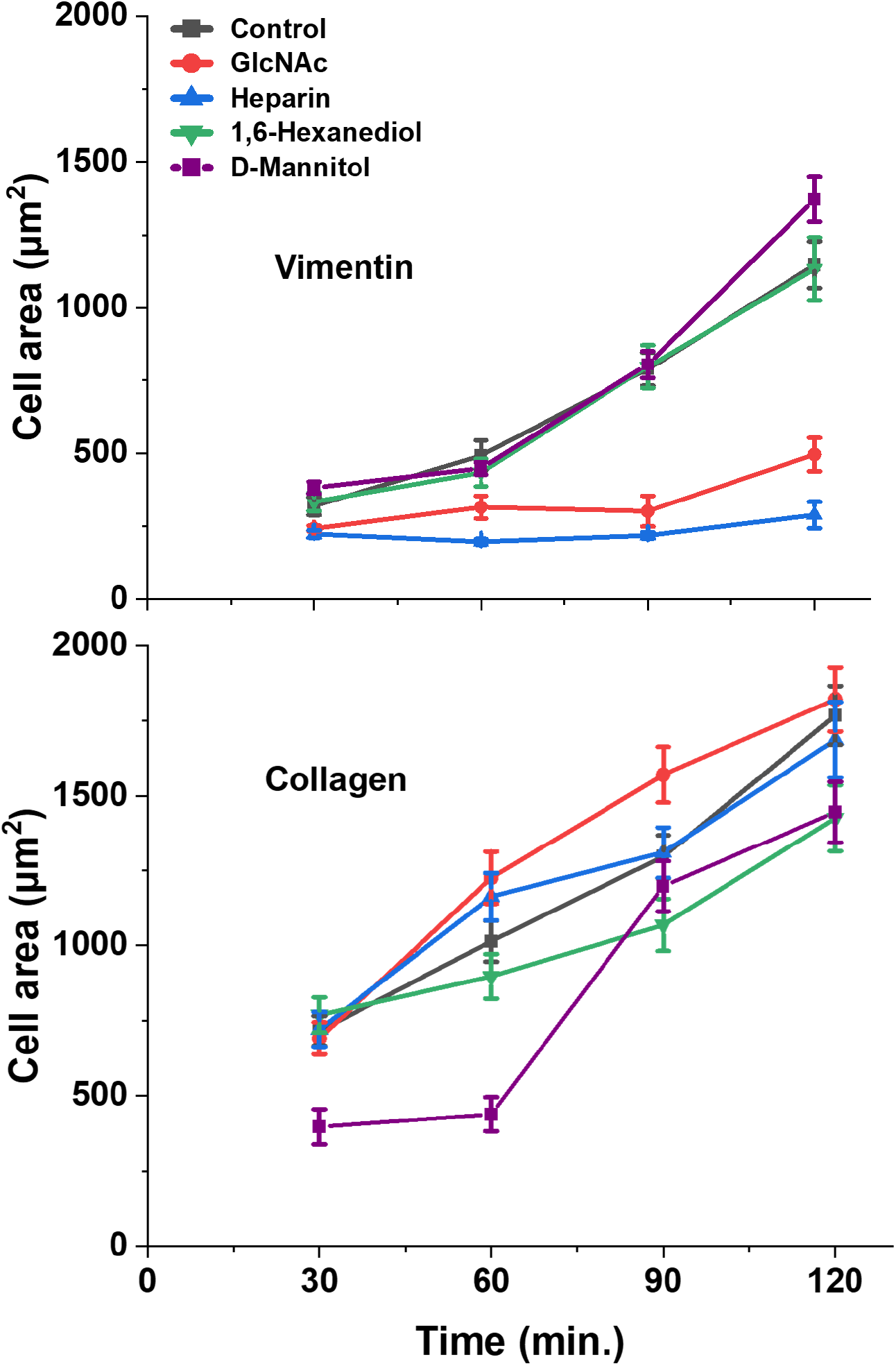
Top panel: Fibroblast spreading is inhibited by GlcNAc and heparin, but not 1,6 hexanediol and D-mannitol on vimentin-coated 30 kPa PAAm gels. Bottom panel: No effect on cells bound to collagen-coated gels.

### Binding of monomeric GlcNAc to vimentin oligomers

To determine if the effects of GlcNAc on cells binding to vimentin-coated gels resulted from direct binding to vimentin, dynamic light scattering (DLS) was done to determine if the apparent size of soluble vimentin oligomers inferred from their diffusion constant was altered by GlcNAc or other solutes. Figure 9 shows that addition of 50 μg/ml GlcNAc to 100 μg/ml soluble vimentin has a large effect on the apparent size of the vimentin oligomer, and heparin sulfate had a small effect in increasing the average hydrodynamic size, mainly by suppressing the numbers of the smallest solutes. Neither D-mannitol or 1,6 hexanediol changed the diffusion time of the vimentin oligomer. Each of the added solutes alone scattered much less light than vimentin oligomers, consistent with their much smaller molecular weight, and did not contribute significantly to the average diffusion constants measured by DLS. Lack of a large effect on the hydrodynamic size of the vimentin oligomer does not rule out possible attachment of small solutes to a single vimentin polymer, but the large effect of GlcNAc is consistent with a capacity to aggregate vimentin oligomers into larger complexes.

**Figure 9.**
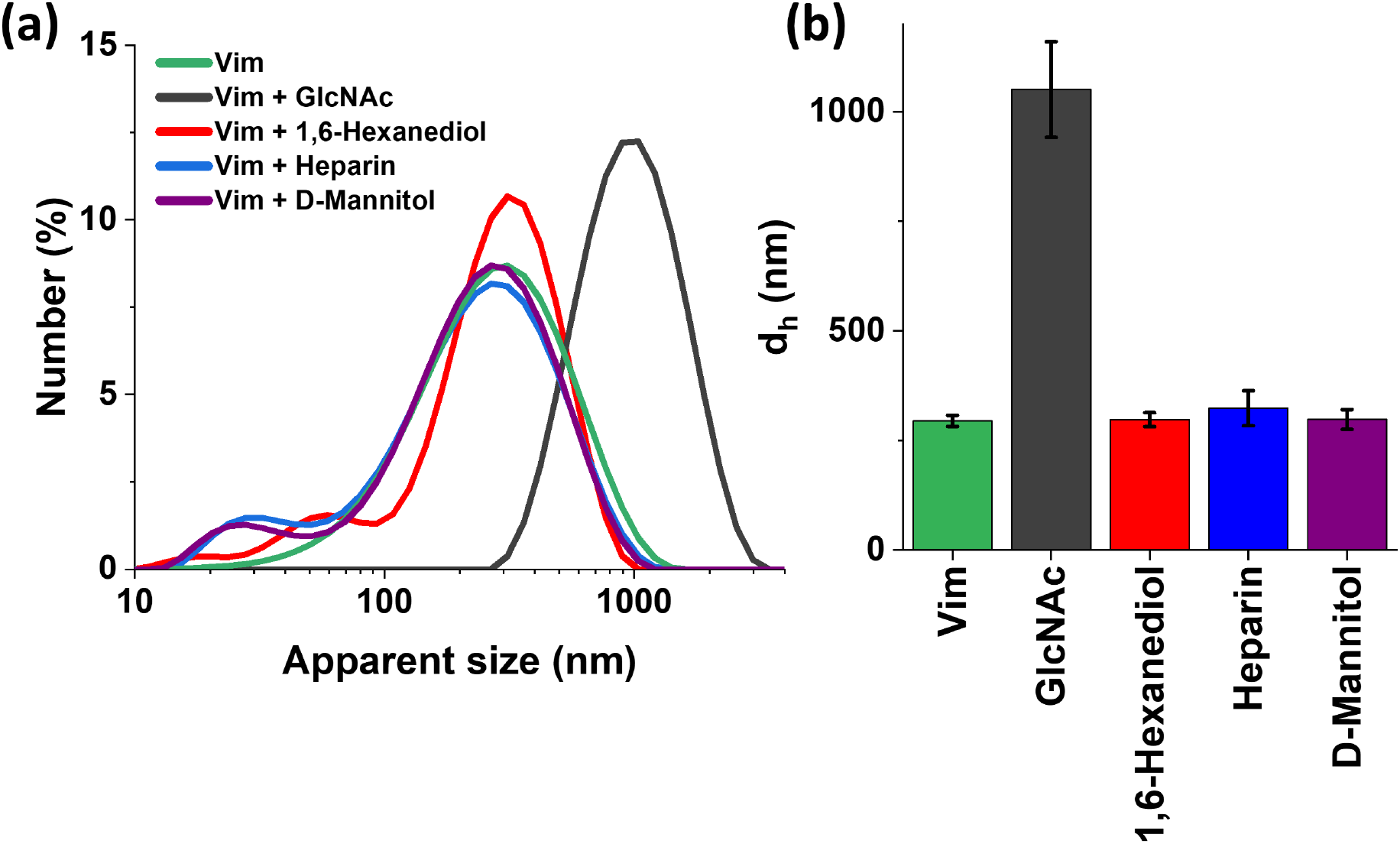
Dynamic light scattering measurement of apparent size of vimentin oligomers after addition of GlcNAc, 1,6 hexanediol, heparin or D-mannitol.

### Effects of soluble vimentin on cell adhesion

Numerous studies have shown that cells release vimentin into the extracellular space in multiple contexts: on the extracellular surface of the cells, attached to the underlying basement membrane, or soluble in the bloodstream or cell culture medium (9,11,13,14). We therefore tested if soluble vimentin had an effect on cell adhesion to substrates that do not already contain vimentin. Figure 10 shows the spreading of fibroblasts on glass when increasing concentrations of vimentin are added in soluble form to the cell culture medium. Both normal and vimentin null fibroblasts are inhibited from spreading on glass by 10 μg/ml concentrations of soluble vimentin, similar to the levels (~1 μg/ml) in serum of patients with cancer or sepsis (47,48). The effect of soluble vimentin appears to be greater on vimentin null fibroblasts compared to normal fibroblasts, suggesting that perhaps the normal fibroblasts already contain some cell surface-bound vimentin, but since the vimentin null fibroblasts do not, they are affected by smaller doses of exogenously added vimentin.

**Figure 10.**
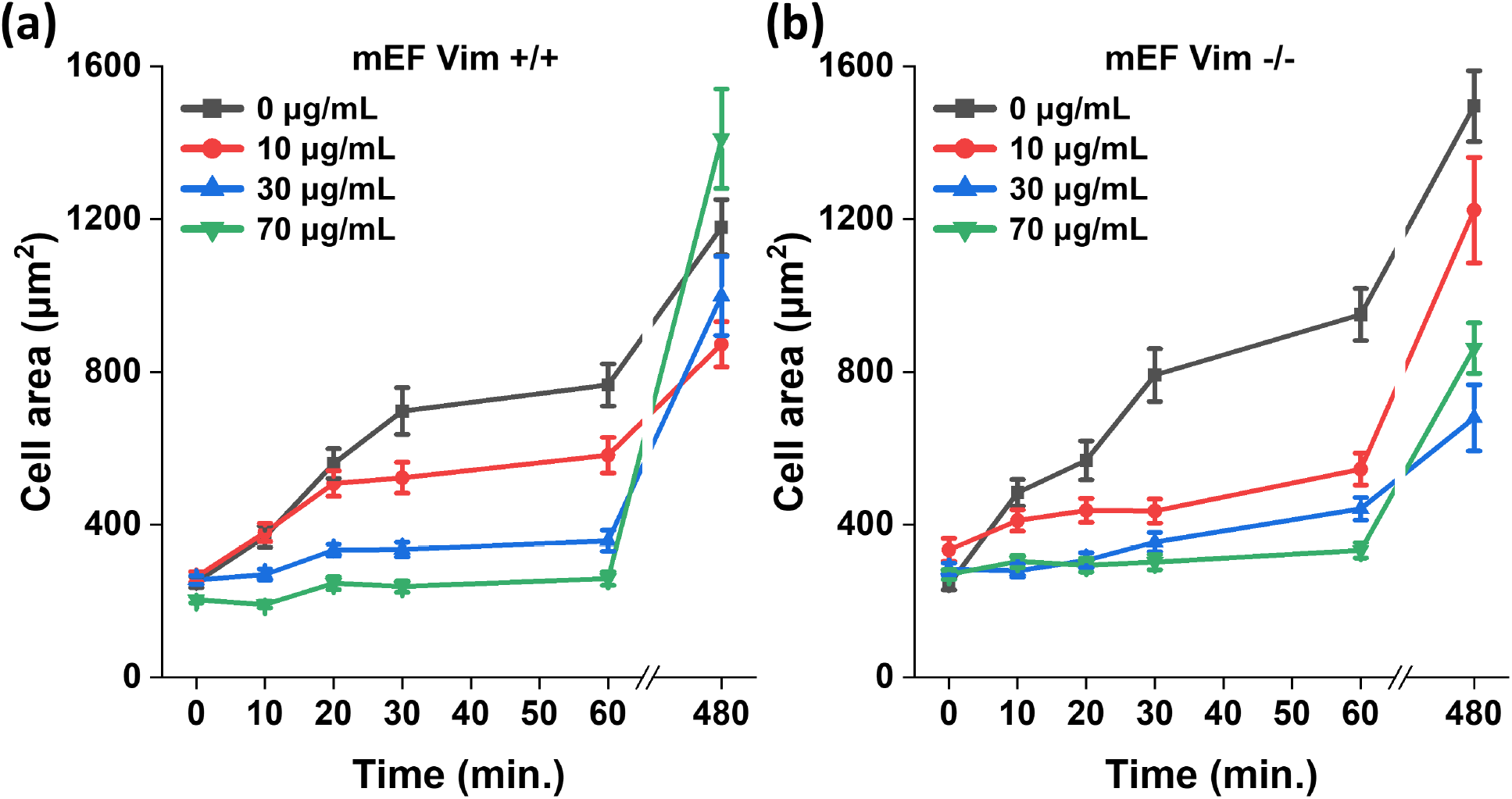
Effect of soluble vimentin on adhesion and spreading of fibroblasts on glass.

## DISCUSSION

Extracellular vimentin is expressed in several pathophysiological conditions, and several reports indicate its role in mediating response to cellular damage or as an autoantigen in rheumatoid arthritis (17,49). Cell surface vimentin interaction with platelets has been implicated in platelet recruitment at injury sites via interaction with von Willebrand Factor (VWF). Pharmacologically disruption, either with anti-vimentin antibodies or with a VWF fragment, positively affected experimental models of ischemic stroke (50). Other reports indicate vimentin interaction with P-selectin, dectin-1, CD44 (51) and insulin-like growth factor 1 receptor (IGF-1R) (52–54). Binding of extracellular vimentin dectin-1, a marker of activated macrophage, triggers reactive oxygen species (ROS) production (54,55). Extracellular vimentin amplifies the secretion of pro-inflammatory cytokines, including IL-6 and TNF-α, induced by oxLDL in macrophages (56). In contrast, LPS-activated dendritic cells, when exposed to vimentin, switch their cytokine profile, decreasing IL-6 and IL-12 secretion, while increasing anti-inflammatory IL-10, thereby reducing Th1 differentiation of naïve T cells (57).

In this study we show that vimentin on an otherwise non-adhesive hydrogel is sufficient for the adhesion of multiple different cell types to the substrate. Binding of cells to vimentin coated substrates appears to involve different receptors from those used to bind collagen or fibronectin because it is inhibited or eliminated either by removal of the cells’ hyaluronic acid-containing glycocalyx, or by addition of soluble N-acetylglucosamine to the substrate, two treatments that have little or no effect on cell binding to collagen- or fibronectin-coated surfaces.

Once bound to vimentin-coated substrates, cells can spread and move, but not divide. Most cell types spread more slowly than they do on collagen-coated surfaces, but in some cases they move more rapidly. The effect of vimentin on solubility is also evident when cell spheroids are placed within three-dimensional collagen networks. The addition of exogenous vimentin, and especially its citrullinated form, causes cells to leave the spheroid and stream out into the collagen network much more efficiently than they would in the absence of exogenous vimentin. These findings confirm the previous reports that extracellular vimentin is an important driver of the movement of fibroblasts into inflamed lung tissue.

The appearance of vimentin in the extracellular space does not appear to be a constitutive aspect of normal physiology, but rather a transient or pathological event that occurs during early development, wound healing, inflammation, and some types of cancer. The nonessential aspect of extracellular vimentin is consistent with the finding that vimentin-null mice develop nearly normally, and are relatively resistant to viral or bacterial infection, or to catastrophic inflammatory responses to endotoxin, two conditions in which extracellular vimentin is normally released onto the cell surface or into the extracellular space (12). Our results here show that vimentin can act as a ligand for cells and suggest that the glycocalyx and glycosylated cell surface proteins that contain N-acetyl glucosamine, serve as adhesion receptors for extracellular vimentin, which has implications for understanding its role in these pathological contexts.

## Data availability

The manuscript contains all data described within the text.

## Acknowledgements

We acknowledge Robert Goldman and Karen Ridge for insightful discussions. This work was supported by NIH R35GM136259 award to PAJ.; NIH R35GM142963 award to AEP, Syracuse University SOURCE award to EW, and the National Science Center of Poland: UMO-2020/01/0/NZ6/00082 awarded to RB.

## Author contributions

RB, DVI, XS, KK, ŁS, KS, JS, and AW performed experiment and analyzed different cell attachment to Vim coated polyacrylamide; PJ, AP and KK performed VIM purification/citrullination; RB, ŁS, and DVI performed immunofluorescence staining; FB performed and analyzed the atomic force microscopy measurements; MT, EW, AP developed the 3D spheroid formation model and analysis; PM and AW contributed to dynamic light scattering experiments analysis. RB, AP, XS, ŁS and PJ wrote the manuscript; PJ, AP, and RB initiated and oversaw the entire project.

## SUPPLEMENTAL DATA

**SI Figure 1.**
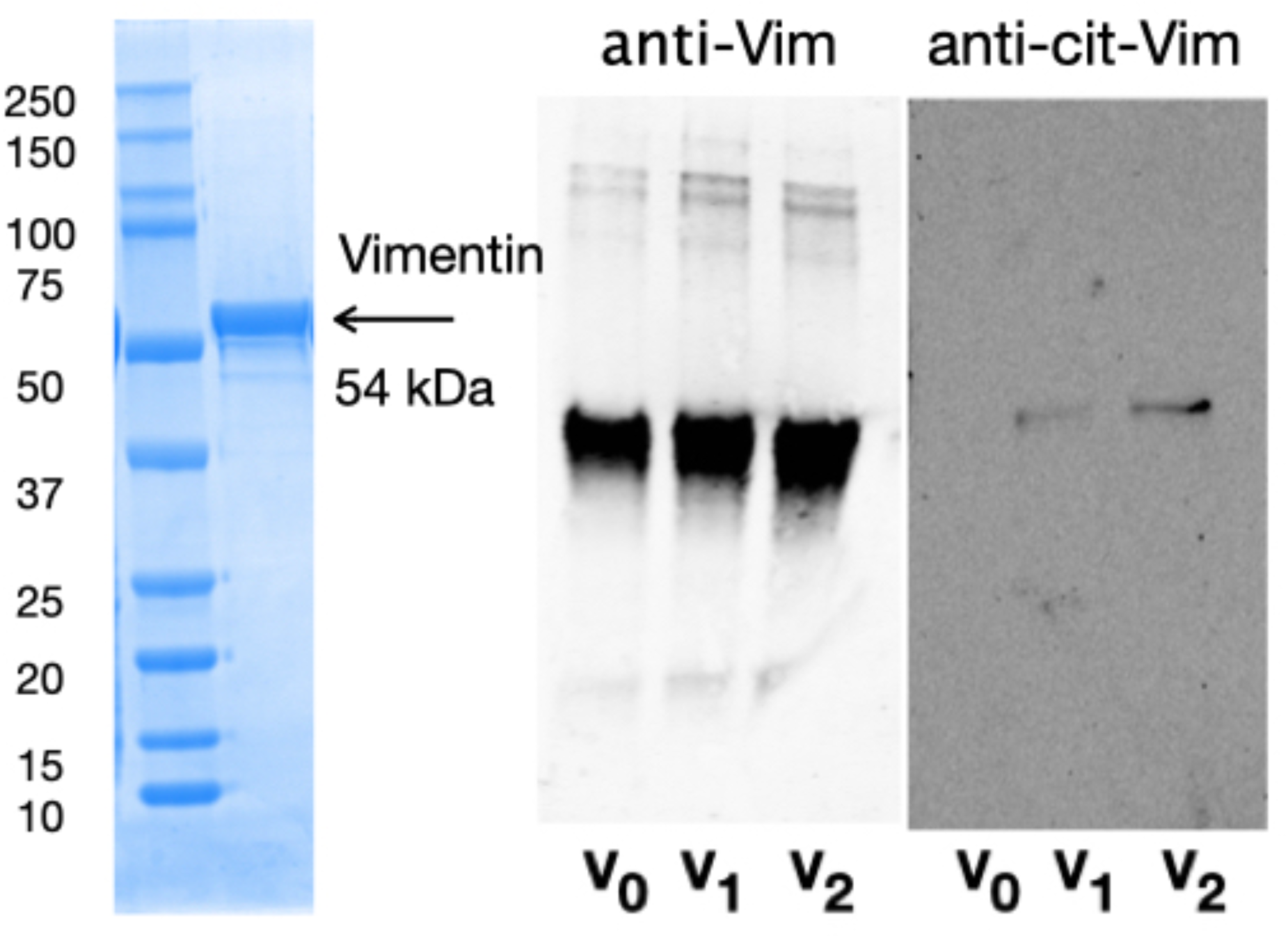
Coomassie-stained SDS-PAGE gel (left) and western blots with anti-vimentin (middle) or anti-citrullinated vimentin (right). v. VO: vimentin; no PAD4. V1: vimentin + PAD4 low salt; V2 vimentin + PAD4 high salt.

**SI Figure 2.**
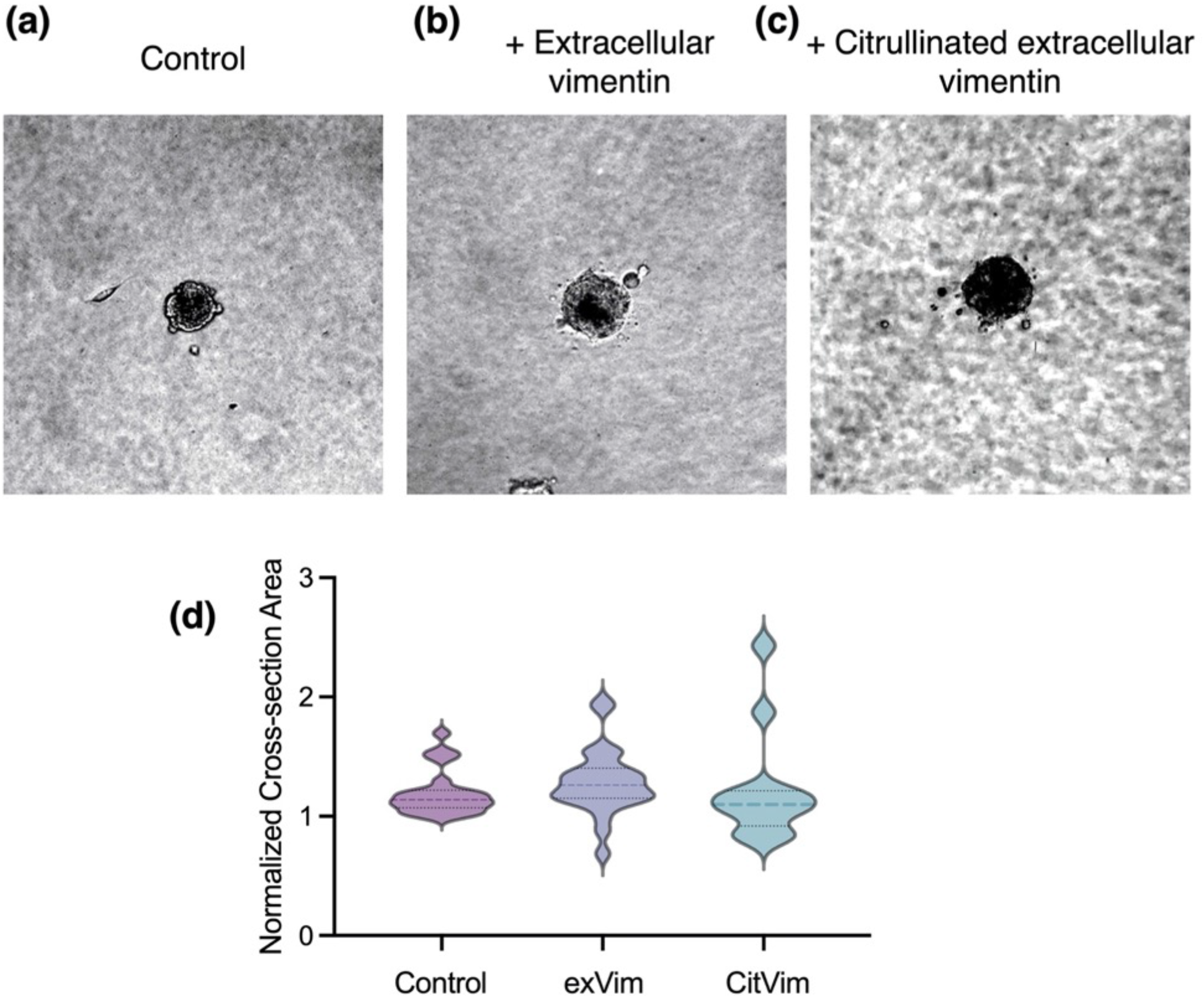
Analysis of vimentin-null cell expansion through collagen matrices with and without extracellular vimentin and its citrullinated form. Representative phasecontrast images of cell aggregates formed by vimentin-null mEF embedded in matrixes comprised of **(a)** collagen I, **(b)** collagen I + vimentin, and **(c)** collagen I + citrullinated vimentin at 24 hours. **(d)** Averaged projected area of cell aggregates formed from vimentin-null mEF embedded in collagen network with vimentin or extracellular vimentin (2 μg/mL).

**SI Figure 3.**
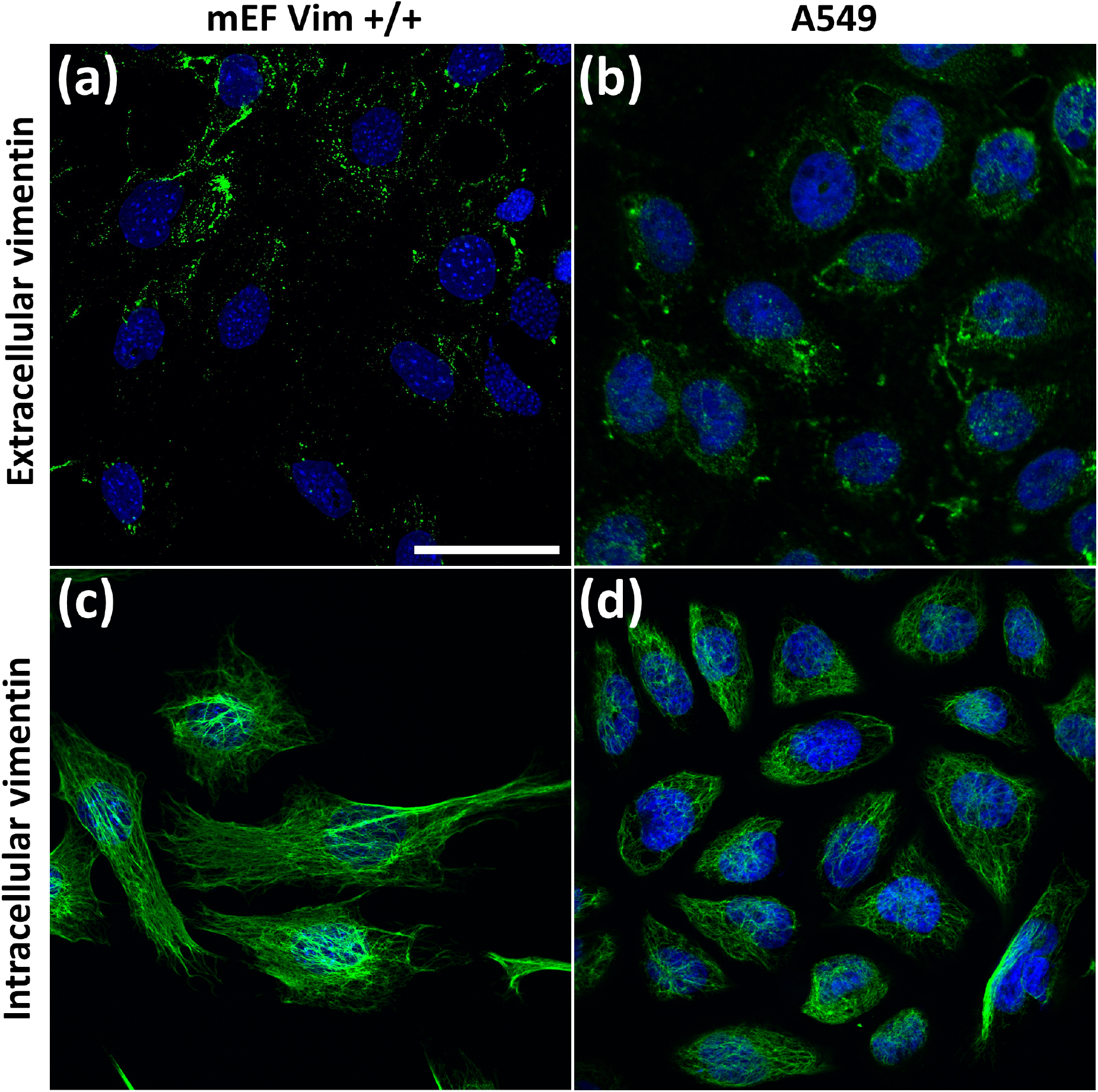
Fluorescence images of extracellular **(a, b)** and intracellular **(c, d)** vimentin in mEF **(a, c)** and A549 **(b, d)** cells. Vimentin is presented in green. Cells were counterstained with DAPI (blue). Scale bar, 50 μm.

## REFERENCES

1. Lind, S. E., Smith, D. B., Janmey, P. A., and Stossel, T. P. (1986) Role of plasma gelsolin and the vitamin D-binding protein in clearing actin from the circulation. J Clin Invest 78, 736–742

2. Janmey, P. A., and Lind, S. E. (1987) Capacity of human serum to depolymerize actin filaments. Blood 70, 524–530

3. Lee, W. M., and Galbraith, R. M. (1992) The Extracellular Actin-Scavenger System and Actin Toxicity. New England Journal of Medicine 326, 1335–1341

4. Pampuscenko, K., Morkuniene, R., Sneideris, T., Smirnovas, V., Budvytyte, R., Valincius, G., Brown, G. C., and Borutaite, V. (2020) Extracellular tau induces microglial phagocytosis of living neurons in cell cultures. J Neurochem 154, 316–329

5. Tobieson, L., Zetterberg, H., Blennow, K., and Marklund, N. (2021) Extracellular fluid, cerebrospinal fluid and plasma biomarkers of axonal and neuronal injury following intracerebral hemorrhage. Sci Rep 11, 16950

6. Ishida, K., Yamada, K., Nishiyama, R., Hashimoto, T., Nishida, I., Abe, Y., Yasui, M., and Iwatsubo, T. (2022) Glymphatic system clears extracellular tau and protects from tau aggregation and neurodegeneration. J Exp Med 219

7. Wei, Y., Liu, M., and Wang, D. (2022) The propagation mechanisms of extracellular tau in Alzheimer’s disease. J Neurol 269, 1164–1181

8. Mor-Vaknin, N., Punturieri, A., Sitwala, K., and Markovitz, D. M. (2003) Vimentin is secreted by activated macrophages. Nat Cell Biol 5, 59–63

9. Danielsson, F., Peterson, M. K., Caldeira Araujo, H., Lautenschlager, F., and Gad, A. K. B. (2018) Vimentin Diversity in Health and Disease. Cells 7

10. Amraei, R., Xia, C., Olejnik, J., White, M. R., Napoleon, M. A., Lotfollahzadeh, S., Hauser, B. M., Schmidt, A. G., Chitalia, V., Muhlberger, E., Costello, C. E., and Rahimi, N. (2022) Extracellular vimentin is an attachment factor that facilitates SARS-CoV-2 entry into human endothelial cells. Proc Natl Acad Sci U S A 119

11. Paulin, D., Lilienbaum, A., Kardjian, S., Agbulut, O., and Li, Z. (2022) Vimentin: Regulation and pathogenesis. Biochimie 197, 96–112

12. Ridge, K. M., Eriksson, J. E., Pekny, M., and Goldman, R. D. (2022) Roles of vimentin in health and disease. Genes Dev 36, 391–407

13. Ramos, I., Stamatakis, K., Oeste, C. L., and Perez-Sala, D. (2020) Vimentin as a Multifaceted Player and Potential Therapeutic Target in Viral Infections. International Journal of Molecular Sciences 21

14. Patteson, A. E., Vahabikashi, A., Goldman, R. D., and Janmey, P. A. (2020) Mechanical and Non-Mechanical Functions of Filamentous and Non-Filamentous Vimentin. Bioessays 42

15. Pogoda, K., Byfield, F., Deptula, P., Ciesluk, M., Suprewicz, L., Sklodowski, K., Shivers, J. L., van Oosten, A., Cruz, K., Tarasovetc, E., Grishchuk, E. L., Mackintosh, F. C., Bucki, R., Patteson, A. E., and Janmey, P. A. (2022) Unique Role of Vimentin Networks in Compression Stiffening of Cells and Protection of Nuclei from Compressive Stress. Nano Lett 22, 4725–4732

16. Block, J., Witt, H., Candelli, A., Danes, J. C., Peterman, E. J., Wuite, G. J., Janshoff, A., and Köster, S. (2018) Viscoelastic properties of vimentin originate from nonequilibrium conformational changes. Science advances 4, eaat1161

17. van Engeland, N. C., Suarez Rodriguez, F., Rivero-Müller, A., Ristori, T., Duran, C. L., Stassen, O. M., Antfolk, D., Driessen, R. C., Ruohonen, S., and Ruohonen, S. T. (2019) Vimentin regulates Notch signaling strength and arterial remodeling in response to hemodynamic stress. Scientific reports 9, 1–14

18. Suprewicz, L., Swoger, M., Gupta, S., Piktel, E., Byfield, F. J., Iwamoto, D. V., Germann, D., Reszec, J., Marcinczyk, N., Carroll, R. J., Lenart, M., Pyre, K., Janmey, P., Schwarz, J. M., Bucki, R., and Patteson, A. (2022) Extracellular Vimentin as a Target Against SARS-CoV-2 Host Cell Invasion. SMALL 18

19. Mor-Vaknin, N., Punturieri, A., Sitwala, K., and Markovitz, D. M. (2003) Vimentin is secreted by activated macrophages. Nature cell biology 5, 59–63

20. Amraei, R., Xia, C., Olejnik, J., White, M. R., Napoleon, M. A., Lotfollahzadeh, S., Hauser, B. M., Schmidt, A. G., Chitalia, V., and Mühlberger, E. (2022) Extracellular vimentin is an attachment factor that facilitates SARS-CoV-2 entry into human endothelial cells. Proceedings of the National Academy of Sciences 119, e2113874119

21. van Beijnum, J. R., Huijbers, E. J. M., van Loon, K., Blanas, A., Akbari, P., Roos, A., Wong, T. J., Denisov, S. S., Hackeng, T. M., Jimenez, C. R., Nowak-Sliwinska, P., and Griffioen, A. W. (2022) Extracellular vimentin mimics VEGF and is a target for anti-angiogenic immunotherapy. Nat Commun 13, 2842

22. Kim, S., Cho, W., Kim, I., Lee, S. H., Oh, G. T., and Park, Y. M. (2020) Oxidized LDL induces vimentin secretion by macrophages and contributes to atherosclerotic inflammation. J Mol Med (Berl)

23. Ise, H., Goto, M., Komura, K., and Akaike, T. (2012) Engulfment and clearance of apoptotic cells based on a GlcNAc-binding lectin-like property of surface vimentin. Glycobiology 22, 788–805

24. Thiam, H. R., Wong, S. L., Qiu, R., Kittisopikul, M., Vahabikashi, A., Goldman, A. E., Goldman, R. D., Wagner, D. D., and Waterman, C. M. (2020) NETosis proceeds by cytoskeleton and endomembrane disassembly and PAD4-mediated chromatin decondensation and nuclear envelope rupture. Proc Natl Acad Sci U S A

25. Khandpur, R., Carmona-Rivera, C., Vivekanandan-Giri, A., Gizinski, A., Yalavarthi, S., Knight, J. S., Friday, S., Li, S., Patel, R. M., Subramanian, V., Thompson, P., Chen, P., Fox, D. A., Pennathur, S., and Kaplan, M. J. (2013) NETs are a source of citrullinated autoantigens and stimulate inflammatory responses in rheumatoid arthritis. Sci Transl Med 5, 178ra140

26. Walker, J. L., Bleaken, B. M., Romisher, A. R., Alnwibit, A. A., and Menko, A. S. (2018) In wound repair vimentin mediates the transition of mesenchymal leader cells to a myofibroblast phenotype. Mol Biol Cell 29, 1555–1570

27. Basta, M. D., Paulson, H., and Walker, J. L. (2021) The local wound environment is a key determinant of the outcome of TGFbeta signaling on the fibrotic response of CD44(+) leader cells in an ex vivo post-cataract-surgery model. Exp Eye Res 213, 108829

28. Parvanian, S., Yan, F., Su, D., Coelho-Rato, L. S., Venu, A. P., Yang, P., Zou, X., Jiu, Y., Chen, H., Eriksson, J. E., and Cheng, F. (2020) Exosomal vimentin from adipocyte progenitors accelerates wound healing. Cytoskeleton (Hoboken) 77, 399–413

29. Adolf, A., Rohrbeck, A., Munster-Wandowski, A., Johansson, M., Kuhn, H. G., Kopp, M. A., Brommer, B., Schwab, J. M., Just, I., Ahnert-Hilger, G., and Holtje, M. (2019) Release of astroglial vimentin by extracellular vesicles: Modulation of binding and internalization of C3 transferase in astrocytes and neurons. Glia 67, 703–717

30. Komura, K., Ise, H., and Akaike, T. (2012) Dynamic behaviors of vimentin induced by interaction with GlcNAc molecules. Glycobiology 22, 1741–1759

31. Griffioen, A. W., Coenen, M. J., Damen, C. A., Hellwig, S. M., HJ van Weering, D., Vooys, W., Blijham, G. H., and Groenewegen, G. (1997) CD44 is involved in tumor angiogenesis; an activation antigen on human endothelial cells. Blood, The Journal of the American Society of Hematology 90, 1150–1159

32. Leterrier, J. F., Kas, J., Hartwig, J., Vegners, R., and Janmey, P. A. (1996) Mechanical effects of neurofilament cross-bridges. Modulation by phosphorylation, lipids, and interactions with F-actin. J Biol Chem 271, 15687–15694

33. van Beers, J. J., Raijmakers, R., Alexander, L. E., Stammen-Vogelzangs, J., Lokate, A. M., Heck, A. J., Schasfoort, R. B., and Pruijn, G. J. (2010) Mapping of citrullinated fibrinogen B-cell epitopes in rheumatoid arthritis by imaging surface plasmon resonance. Arthritis Res Ther 12, R219

34. Inagaki, M., Takahara, H., Nishi, Y., Sugawara, K., and Sato, C. (1989) Ca2+-dependent deimination-induced disassembly of intermediate filaments involves specific modification of the amino-terminal head domain. J Biol Chem 264, 18119–18127

35. Kandow, C. E., Georges, P. C., Janmey, P. A., and Beningo, K. A. (2007) Polyacrylamide hydrogels for cell mechanics: steps toward optimization and alternative uses. Methods Cell Biol 83, 29–46

36. Pogoda, K., Charrier, E. E., and Janmey, P. A. (2021) A Novel Method to Make Polyacrylamide Gels with Mechanical Properties Resembling those of Biological Tissues. Bio Protoc 11, e4131

37. Mandal, K., Raz-Ben Aroush, D., Graber, Z. T., Wu, B., Park, C. Y., Fredberg, J. J., Guo, W., Baumgart, T., and Janmey, P. A. (2019) Soft Hyaluronic Gels Promote Cell Spreading, Stress Fibers, Focal Adhesion, and Membrane Tension by Phosphoinositide Signaling, Not Traction Force. ACS Nano 13, 203–214

38. Zancla, A., Mozetic, P., Orsini, M., Forte, G., and Rainer, A. (2022) A primer to traction force microscopy. J Biol Chem 298, 101867

39. https://ibidi.com/img/cms/support/AN/AN32_Generation_of_spheroids.pdf.

40. https://ibidi.com/img/cms/support/AN/AN26_CollagenI_protocols.pdf.

41. Li, F. J., Surolia, R., Li, H., Wang, Z., Liu, G., Kulkarni, T., Massicano, A. V. F., Mobley, J. A., Mondal, S., de Andrade, J. A., Coonrod, S. A., Thompson, P. R., Wille, K., Lapi, S. E., Athar, M., Thannickal, V. J., Carter, A. B., and Antony, V. B. (2021) Citrullinated vimentin mediates development and progression of lung fibrosis. Science Translational Medicine 13

42. Menko, A. S., Romisher, A., and Walker, J. L. (2022) The Pro-fibrotic Response of Mesenchymal Leader Cells to Lens Wounding Involves Hyaluronic Acid, Its Receptor RHAMM, and Vimentin. Front Cell Dev Biol 10, 862423

43. Pan, Y., Lei, T., Teng, B., Liu, J., Zhang, J., An, Y., Xiao, Y., Han, J., Pan, X., Wang, J., Yu, H., Ren, H., and Li, X. (2011) Role of vimentin in the inhibitory effects of low-molecular-weight heparin on PC-3M cell adhesion to, and migration through, endothelium. J Pharmacol Exp Ther 339, 82–92

44. Paszek, M. J., Boettiger, D., Weaver, V. M., and Hammer, D. A. (2009) Integrin clustering is driven by mechanical resistance from the glycocalyx and the substrate. PLoS Comput Biol 5, e1000604

45. Ise, H., Kobayashi, S., Goto, M., Sato, T., Kawakubo, M., Takahashi, M., Ikeda, U., and Akaike, T. (2010) Vimentin and desmin possess GlcNAc-binding lectin-like properties on cell surfaces. Glycobiology 20, 843–864

46. Zhou, X., Lin, Y., Kato, M., Mori, E., Liszczak, G., Sutherland, L., Sysoev, V. O., Murray, D. T., Tycko, R., and McKnight, S. L. (2021) Transiently structured head domains control intermediate filament assembly. Proc Natl Acad Sci U S A 118

47. Arko-Boham, B., Lomotey, J. T., Tetteh, E. N., Tagoe, E. A., Aryee, N. A., Owusu, E. A., Okai, I., Blay, R. M., and Clegg-Lamptey, J. N. (2017) Higher serum concentrations of vimentin and DAKP1 are associated with aggressive breast tumour phenotypes in Ghanaian women. Biomark Res 5, 21

48. Su, L., Pan, P., Yan, P., Long, Y., Zhou, X., Wang, X., Zhou, R., Wen, B., Xie, L., and Liu, D. (2019) Role of vimentin in modulating immune cell apoptosis and inflammatory responses in sepsis. Sci Rep 9, 5747

49. Van Steendam, K., Tilleman, K., and Deforce, D. (2011) The relevance of citrullinated vimentin in the production of antibodies against citrullinated proteins and the pathogenesis of rheumatoid arthritis. Rheumatology 50, 830–837

50. Fasipe, T. A., Hong, S.-H., Da, Q., Valladolid, C., Lahey, M. T., Richards, L. M., Dunn, A. K., Cruz, M. A., and Marrelli, S. P. (2018) Extracellular vimentin/VWF (von Willebrand factor) interaction contributes to VWF string formation and stroke pathology. Stroke 49, 2536–2540

51. Päll, T., Pink, A., Kasak, L., Turkina, M., Anderson, W., Valkna, A., and Kogerman, P. (2011) Soluble CD44 interacts with intermediate filament protein vimentin on endothelial cell surface. PloS one 6, e29305

52. Lam, F. W., Da, Q., Guillory, B., and Cruz, M. A. (2018) Recombinant human vimentin binds to P-selectin and blocks neutrophil capture and rolling on platelets and endothelium. The Journal of Immunology 200, 1718–1726

53. Shigyo, M., Kuboyama, T., Sawai, Y., Tada-Umezaki, M., and Tohda, C. (2015) Extracellular vimentin interacts with insulin-like growth factor 1 receptor to promote axonal growth. Scientific reports 5, 1–13

54. Thiagarajan, P. S., Yakubenko, V. P., Elsori, D. H., Yadav, S. P., Willard, B., Tan, C. D., Rene Rodriguez, E., Febbraio, M., and Cathcart, M. K. (2013) Vimentin is an endogenous ligand for the pattern recognition receptor Dectin-1. Cardiovascular research 99, 494–504

55. Nandakumar, V., Hebrink, D., Jenson, P., Kottom, T., and Limper, A. H. (2017) Differential macrophage polarization from pneumocystis in immunocompetent and immunosuppressed hosts: potential adjunctive therapy during pneumonia. Infection and immunity 85, e00939-00916

56. Kim, S., Cho, W., Kim, I., Lee, S.-H., Oh, G. T., and Park, Y. M. (2020) Oxidized LDL induces vimentin secretion by macrophages and contributes to atherosclerotic inflammation. Journal of Molecular Medicine 98, 973–983

57. Yu, M. B., Guerra, J., Firek, A., and Langridge, W. H. (2018) Extracellular vimentin modulates human dendritic cell activation. Molecular immunology 104, 37–46

